# Automated design of synthetic microbial communities

**DOI:** 10.1101/2020.06.30.180281

**Authors:** Behzad D. Karkaria, Alex J. H. Fedorec, Chris P. Barnes

## Abstract

In naturally occurring microbial systems, species rarely exist in isolation. There is strong ecological evidence for a positive relationship between species diversity and the functional output of communities. The pervasiveness of these communities in nature highlights that there may be advantages for engineered strains to exist in cocultures as well. Building synthetic microbial communities allows us to create distributed systems that mitigates issues often found in engineering a monoculture, especially when functional complexity is increasing. Here, we demonstrate a methodology for designing robust synthetic communities that use quorum sensing to control amensal bacteriocin interactions in a chemostat environment. We explore model spaces for two and three strain systems, using Bayesian methods to perform model selection, and identify the most robust candidates for producing stable steady state communities. Our findings highlight important interaction motifs that provide stability, and identify requirements for selecting genetic parts and tuning the community composition.

## Introduction

Traditionally, in biotechnology and synthetic biology, a microbe is engineered and grown as a monoculture to perform a particular function. Novel functionality is imparted by introducing heterologous genetic processes that would not normally be found in the organism. Unintended interactions between the introduced heterologous processes can cause the engineered function to behave in an unintended manner [1, 2, 3]. Metabolic burden imposed by heterologous expression can significantly slow growth rates and encourage selection of mutants [4]. The limited cellular resource availability can cause circuits to behave differently when expressed alongside one another [5, 6]. Opportunities for unintended interactions will increase exponentially with the number of heterologous processes introduced into a host strain. These limitations are potentially circumvented with the use of microbial communities, which can be used to implement distributed systems. A distributed system would allow us to allocate functional components between subpopulations of cells, creating physical barriers that insulate processes from one another and distributing the burden of heterologous expression [7]. This allows us to scale complexity in a manner that could not be achieved under the limitations of a monoculture. In natural environments, we observe mixed-species microbial communities that exhibit competitive advantages over monocultures in productivity, resource efficiency, metabolic complexity and resistance to invasion [8, 9]. Being able to predictably and reproducibly construct microbial communities for synthetic biology or biotechnology applications would allow us to harness these advantages.

While the potential advantages of building distributed systems for synthetic biology applications are clear, the maintenance and control of microbial communities comes with its own challenges. The competitive exclusion principle states that when multiple populations compete for a single limiting resource (in the absence of other interactions), a single population with the highest fitness will drive the others to extinction [10]. Evidence from natural microbial systems and ecological studies have shown us that stability can arise through feedback between subpopulations. Both cooperative and competitive interactions are important for integrating feedback that can stabilise communities by manipulating growth or fitness of the subpopulations [11, 12, 13, 14, 15, 16].

Communication between members of a microbial community is important for the dynamic regulation of its behaviour. Quorum sensing (QS) systems are a key set of tools that enable us to engineer communication between and within subpopulations of a community. In natural systems, they can be used to regulate genetic processes as a function of population density [17]. QS systems consist of a biosynthetic module that produces small, freely diffusible molecules. These molecules bind regulatory proteins that can activate or repress gene expression at specific promoters [17]. Synthetic microbial communities have been built using QS to regulate processes that manipulate the growth rate or fitness of a population. Fitness can be manipulated by the expression of lysis proteins, metabolic enzymes, toxins and anti-microbial peptides (AMPs) [14, 18, 19, 20, 21, 22, 23, 24]. In this study we focus on exploring the use of bacteriocins to manipulate subpopulation growth rates. Bacteriocins are gene-encoded AMPs that can be used to directly suppress the growth rate of a sensitive population [25]. They are exported into the extracellular environment, and generally use “Trojan horse” strategies to enter and kill sensitive strains. One of the most well studied bacteriocins is microcin V (MccV), a broad spectrum class IIa bacteriocin [25]. It is exported by the cell, and takes advantage of iron-binding receptors displayed extracellularly to enter a sensitive cell, having an amensal effect on the target [25]. Expression of immunity genes provide protection against the bacteriocin and these can be expressed separately or in conjunction with the bacteriocin [26]. Bacteriocins present themselves as a versatile actuator of population growth. Their expression by a single strain can affect any number of subpopulations in a synthetic community. This is in contrast to lysis proteins and intracellular toxins which affect individual cells. Previously, we have demonstrated the use of MccV expression to improve plasmid maintenance in a population [27] and for building stable cocultures that overcome competitive exclusion [28]. Other bacteriocins, such as nisin, have also been used to produce stable communities [24].

Microbial communities exhibit diverse temporal dynamics including stable steady state, oscillations and deterministic chaos [29, 30]. An important part of designing synthetic communities is being able to define the temporal dynamics that suit the intended application. In this work we set an objective of producing robust stable steady state communities in a chemostat environment. Stable communities allow multiple subpopulations to contribute in a specialised manner to an aggregate community function. We envisage that through the expression of QS and bacteriocins we can build ‘plug-and-play’ stabilising systems. These consist of a set of genetic parts that when expressed by the appropriate subpopulations will produce a stable steady state community. The design challenge lies in identifying the most robust combinations of QS and bacteriocin expression, giving us the highest probability of producing stable steady state in a chemostat environment. Given a small set of genetic parts, the number of ways they can be combined to produce a candidate system can become overwhelming; system design by intuition alone becomes increasingly challenging when dealing with multi-level interactions. Predicting how a system will behave before implementation is essential for the efficient use of lab resources and fully understanding the interactions that occur [31]. Models allow us to make data driven decisions concerning wet lab implementation. Model selection is a process of identifying good performing models from a selection of candidates [32]. Approximate Bayesian computation with sequential Monte Carlo sampling (ABC SMC) is a model selection and parameterisation method that has been applied to synthetic biology systems [33]. We have previously used ABC SMC to identify robust genetic oscillators in two and three-gene networks [34], and to identify parameter regions that give rise to multistable genetic switches [35]. ABC SMC also allows us to derive design principles that are key to producing the desired behaviour [34, 35]. Similarly, Yeoh et al. developed software that compared the ability of genetic parts to produce logic gate behaviours, performing model selection using Akaike information criterion (AIC) [36]. Workflows have been developed to design regulatory networks from databases of characterised parts with defined behaviours [37, 38]. Automated circuit design has the potential to greatly improve the engineering process in synthetic biology.

Here, we present the first instance of automated synthetic community design. Our workflow automatically generates candidate systems from a set of parts which can be used to engineer a community. We use ABC SMC to perform model selection, identifying candidate systems that have the highest probability of producing stable communities in a chemostat bioreactor. Using these methods we reveal the optimal designs for two- and three-strain systems. This workflow also allows us to derive fundamental design principles for building stable communities and reveal critical parameters to control the community composition.

## Results

### Automated synthetic microbial Community Designer (AutoCD) workflow

Figure 1 illustrates AutoCD, the workflow developed and applied in this study. First, we set the available parts which can be used to build a stabilising system in a chemostat environment. This consists of the number of strains (*N*), bacteriocins (*B*), and QS systems (*A*). Any *A* in a system can regulate the expression of any *B* in the system by induction or repression. Strains in all models are dependent upon a single nutrient resource (*S*), which is consumed by strains and replenished through dilution of the chemostat with fresh media. Importantly, all models therefore include nutrient based competition between subpopulations. Uniform prior distributions are set, describing each part and their interactions with one another (Table 2). The uniform priors we use span the expected ranges these parameters could take. Importantly, in scenarios where the particular parts have already been selected and characterised, the prior parameters can be constrained. The available parts and prior parameter distributions serve as inputs to the Model Space Generator, which conducts a series of combinatorial steps to produce all possible genetic circuits. The Model Space Generator then builds unique combinations of strains expressing different genetic circuits, where each combination is a candidate stabilising system. Filtering steps remove inviable, redundant and mirror systems, yielding a set of unique candidates to be assessed (*Methods*). The Model Space Generator produces an ordinary differential equation (ODE) model for each system in the context of the chemostat environment, and these models form our prior model space.

**Table 1:** Bayes factor categorisation to describe evidence in favour of *m*_1_, compared with *m*_2_.

| Bayes factor ($BF$)<br>value | Evidence against $m_2$<br>(in favour of $m_1$) |
| --- | --- |
| 1 to 3 | Very weak |
| 3 to 20 | Positive |
| 20 to 150 | Strong |
| > 150 | Very strong |

**Table 2:** Uniform priors used to for parameter sampling and initial species values.

| Parameter / Species | Description | Prior (min) | Prior (max) | Units | Citation |
| --- | --- | --- | --- | --- | --- |
| Parameters |  |  |  |  |  |
| $C_N$ | OD to cell number scaling factor | 1e9 | 1e9 | None | N/A |
| $C_B$ | Microcin scaling factor | 1e-9 | 1e-9 | None | N/A |
| $C_A$ | QS scaling factor | 1e-9 | 1e-9 | None | N/A |
| $D$ | Dilution rate | 0.01 | 0.2 | $h^{-1}$ | N/A |
| $K_{A_y B_z}$ | Half maximal QS promoter activation/repression from $A_y$ to $B_z$ | 1e-9 | 1e-6 | $M$ | [65] |
| $K_x$ | Monod's half saturation constant | 3.9e-5 | 3.9e-5 | $M$ | [66] |
| $K_\omega$ | Half saturation killing constant | 1e-7 | 1e-6 | $M$ | [67, 68] |
| $S_0$ | Substrate concentration of input media (0.4% glucose) | 0.02 | 0.02 | $M$ | M9 media |
| $\gamma$ | E. coli substrate yield | 1e11 | 1e11 | cell $M^{-1}$ | [69] |
| $k_{A_y}$ | Production rate of AHL per cell | 1e-22 | 1e-15 | $M h^{-1}$ | [70] |
| $KB_{max} z$ | Maximal expression rate of microcin | 1e-22 | 1e-15 | $M h^{-1}$ | [6] |
| $\mu_{x_{max}}$ | Maximum growth rate | 0.4 | 3 | $h^{-1}$ | [42, 43] |
| $n_z$ | Hill coefficient AHL induced expression | 1 | 2 | $M$ | [65] |
| $n_\omega$ | Hill coefficient for killing | 1 | 2 | $M$ | [65] |
| $\omega_{max}$ | Maximum rate of bacteriocin killing | 0.5 | 2.0 | $M^{-1} h^{-1}$ | [71, 67, 68] |
| Initial species |  |  |  |  |  |
| $N$ | OD of strain | 0.01 | 0.5 | OD | N/A |
| $S$ | 0.4% glucose concentration | 0.02 | 0.02 | $M$ | N/A |
| $B$ | Microcin concentration | 1e-81 | 1e-81 | $M$ | N/A |
| $A$ | QS concentration | 1e-10 | 1e-10 | $M$ | N/A |

**Figure 1:**
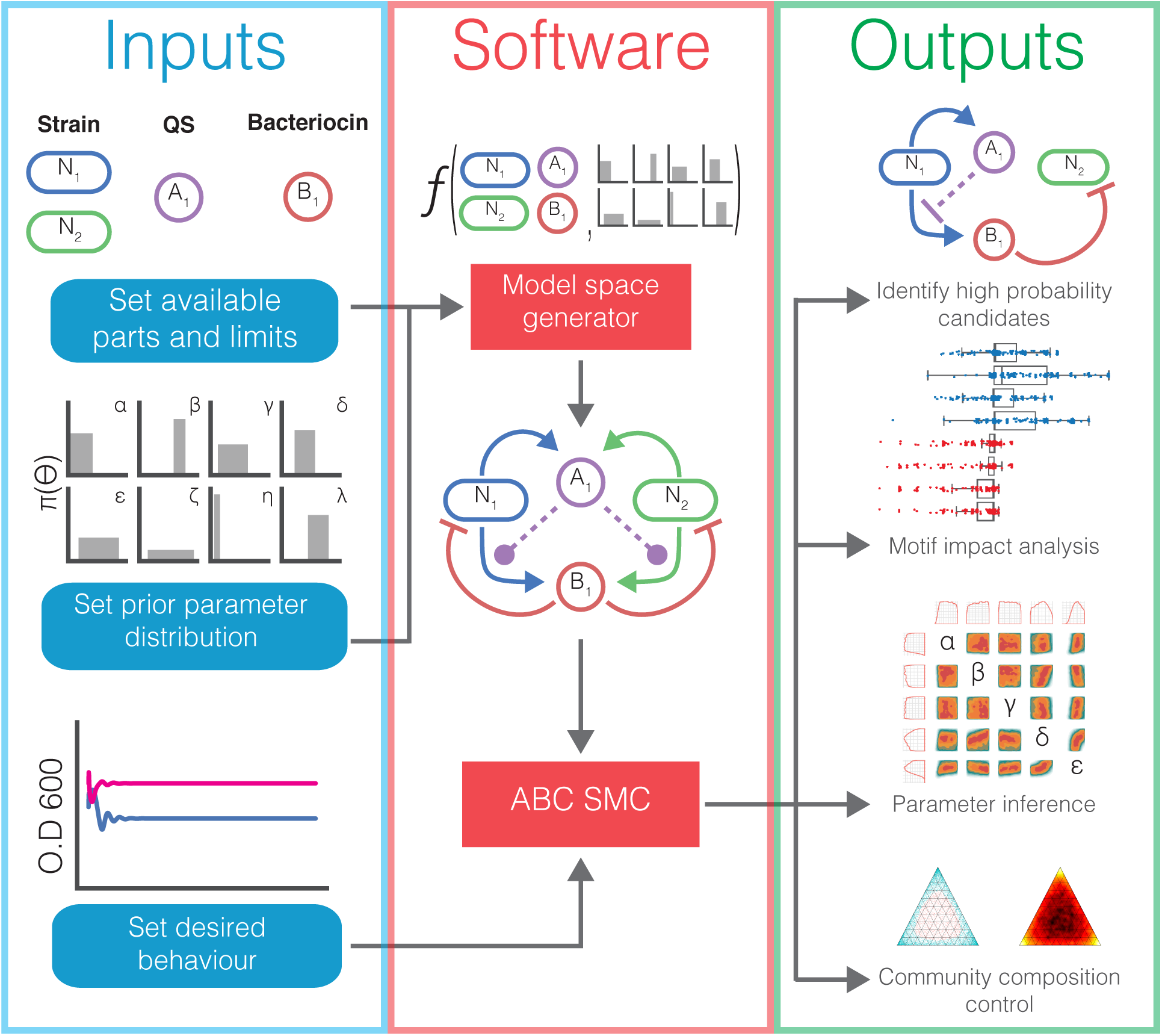
Overview of AutoCD pipeline. Model selection workflow begins with definition of available parts and prior parameter distributions, used to generate system models from all the possible interactions. We use ABC SMC to perform model selection for the desired population behaviour. The outputs of ABC SMC provide us with community designs, insight into underlying motifs, parameter requirements and information on composition tunability.

The final required input is a mathematical description of the objective population behaviour, a stable steady state. We use three distance functions (*d*_1_, *d*_2_, *d*_3_) to describe how far away a simulation is from the objective stable steady state (Eq 1). *d*_1_ is the final gradient of *N*_*x*_, capturing the most fundamental characteristic of stable steady state where the population level of a strain is unchanging. *d*_2_ is the standard deviation of a population, chosen to detect unstable behaviours such as oscillations, favouring simulations that reach stable steady state quickly. *d*_3_ is the reciprocal of the strain population at the end of the simulation, allowing us to define a minimum population density. Given the three distances, *ϵ*_*F*_ defines thresholds below which a simulation meets the requirements of our stable steady state objective. The distances of all strain populations in a simulation must be below these thresholds to satisfy the objective behaviour. 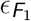 was chosen to match the error tolerance of the ODE solver and 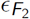 threshold was chosen through qualitative assessment of simulation data to define a practical threshold for what stable steady state simulations should look like. 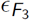 is set to ensure all populations have a minimum final *OD* of 0.001. The posterior distribution is made up of simulations where the distances for each strain population are less than the *ϵ*_*F*_ thresholds (Eq 2). ABC SMC performs model selection on the model space for the objective defined by these distance functions and *ϵ*_*F*_. A particle is a sampled model and sampled parameters. ABC SMC initially samples particles from the prior distributions with an unbounded distance threshold. Particles are propagated through intermediate distributions, gradually reducing the distance thresholds until they equal *ϵ*_*F*_ (*Methods*). ABC SMC provides an estimation of model and parameter space posterior probabilities for the given prior distributions and the objective behaviour. The output of ABC SMC is an approximation of the posterior distribution which we can use to help us design synthetic communities and chemostat settings in the lab.

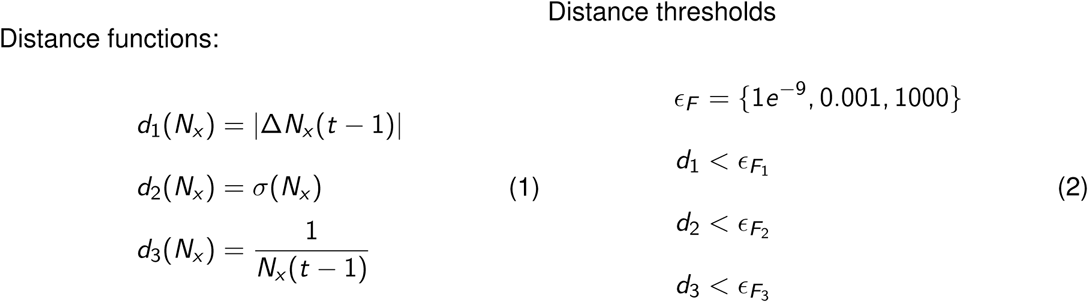

### Designing two strain cocultures that achieve steady state

Here we apply AutoCD to the design of a stable steady state coculture containing two strains. In Figure 2 we define a model space consisting of two strains (*N*_1_, *N*_2_), two bacteriocins (*B*_1_, *B*_2_) and two QS systems (*A*_1_, *A*_2_). We set model space limits to enable feasible experimental implementation, allowing expression of up to one QS per strain and expression of up to one bacteriocin per strain. Each strain can be sensitive to up to one bacteriocin. Given these conditions, the Model Space Generator yields 69 unique two strain models (*m*_0_, *m*_1_…*m*_68_). These 69 models serve as a uniform prior model space upon which we perform model selection using ABC SMC. From the available genetic parts, there are 17 possible interaction options that could exist between species in each candidate model. We perform hierarchical clustering on the interactions present in each model, grouping models based on the similarity of their interactions. This clustering is visualised as a dendrogram in (Figure 2A). ABC SMC approximates the posterior probability of each model for the stable steady state objective, indicating how effective the candidate system is in producing stable steady state. *m*_62_ has the highest posterior probability, and is therefore the system which most robustly produces stable steady state (Figure 2A). *m*_62_ consists of two strains exhibiting a cross-protection mutualism relationship [30]. Each strain expresses an orthogonal QS molecule that represses the expression of a self-limiting bacteriocin in the opposing strain (Figure 2B). This is an interdependence between the two strains where the extinction of one strain would result in the extinction of the other. The exclusion of one strain reduces the fitness of the other, creating a closed feedback loop which overcomes the competitive exclusion principle.

**Figure 2:**
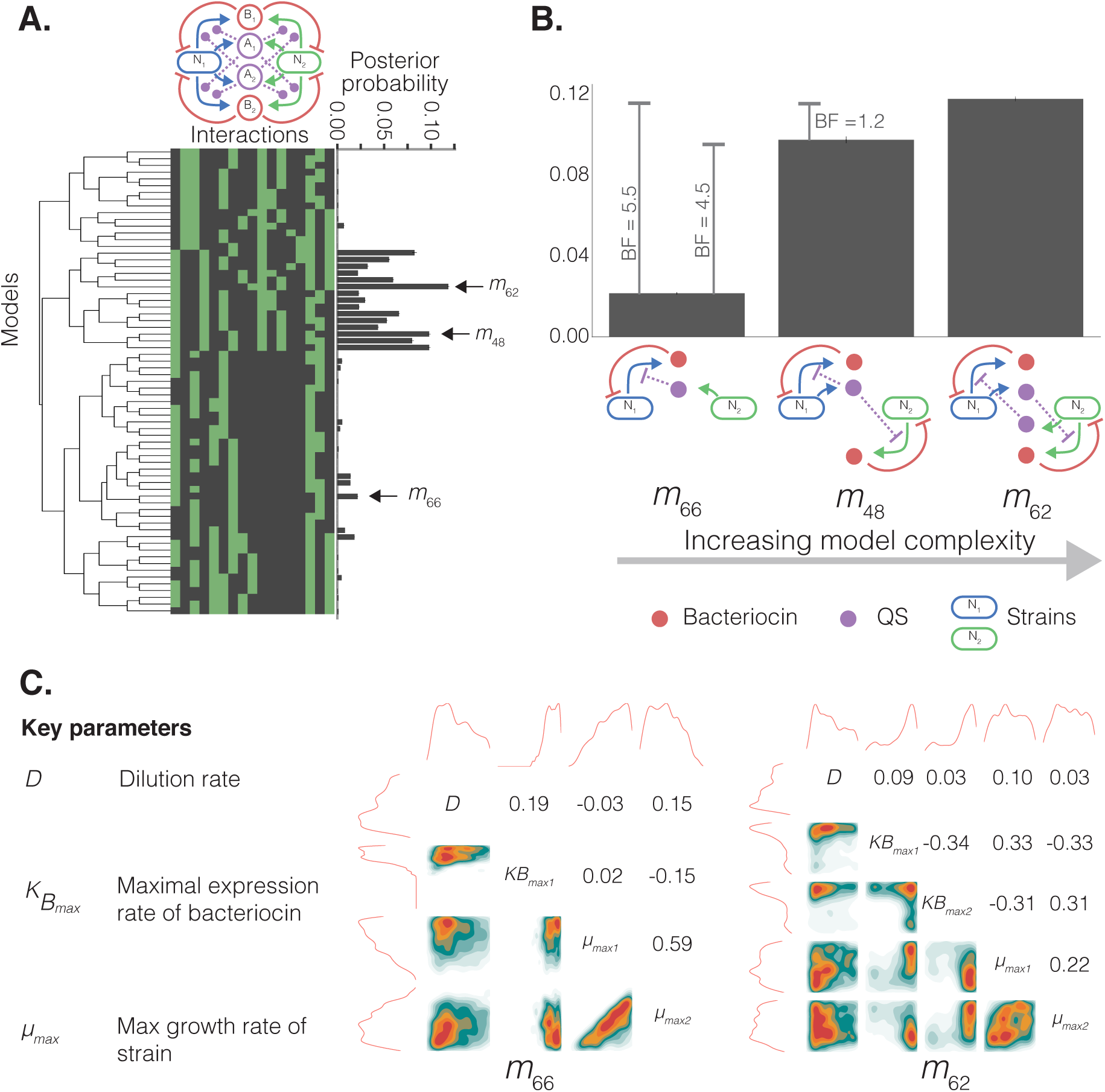
Output of AutoCD for the two strain stable steady state objective A.) Dendrogram is generated by hierarchical clustering of the adjacency matrices for each model in the two strain model space. All possible interactions are shown in the network illustration. Each column of the heatmap represents a possible interaction between species, where green indicates the interaction exists for the model and black indicates absence of the interaction. The bar chart shows the posterior probability of each model. B) Shows the models with highest posterior probability when subsetted for number of parts expressed, in order of increasing complexity (2, 3 and 4 parts). Bayes factors (BF) are shown for pairwise comparison of the three models and error bars show the standard deviation across three repeat experiments. Model schematics show the interactions between strains (blue and green), bacteriocin (red) and QS molecules (purple). C) Posterior parameter distributions of several tunable parameters in *m*_66_ and *m*_62_. The top and left plots show 1D density distributions of each parameter, central distributions are 2D density distributions for each pair of parameters. Pearson correlation coefficients are shown on the top side of the diagonal for each parameter pair.

When designing new systems, minimising the number of genetic parts will reduce the number of experimental variables, improving the ease of construction and optimisation of a system. We subset the model space by the number of expressed species in the system (maximum two QS and two bacteriocin), yielding subsets containing candidate models with two, three and four expressed species (low complexity to high complexity). We identify the candidates with the highest posterior probability in each subset (Figure 2B). We see the posterior probability increasing despite the larger parameter spaces, which is important because ABC SMC will naturally favour models which yield stable steady state with the smallest possible number of parameters (Occam’s razor) [39]. We see that all three models have self-limiting motifs, where a strain is sensitive to the bacteriocin it produces. All three models are devoid of other-limiting motifs, where a strain is sensitive to a bacteriocin produced by another strain.

The Bayes Factor (*BF*) is a ratio between the marginal likelihoods of two models, giving a quantification of support for one model compared with another. *BF* > 3.0 indicates evidence of a notable difference between the two models, while *BF* < 3.0 suggests insubstantial evidence [40] (Table 1). The *BF* of *m*_66_ compared with *m*_48_ suggests substantial improvement in the posterior probability can be made by increasing complexity. However, the *BF* of *m*_48_ compared with *m*_62_ suggests insubstantial evidence behind this improvement in posterior probability. These diminishing returns when increasing system complexity hold important ramifications for system design. The introduction of an additional QS part to move from *m*_48_ to *m*_62_ may not be efficient for the minor improvement in steady state robustness.

Model selection has identified the best performing designs for producing stable communities. However, the parts used in the design may require specific characteristics or chemostat experimental setup. ABC SMC also produces posterior parameter distributions for each model, giving us information about the parameter values necessary to yield stable steady state. Figure 2C shows the posterior distributions of several tunable parameters in *m*_66_ and *m*_62_, showing the parameter values that give rise to stable communities. The dilution rate of the chemostat (*D*) is a directly tunable parameter and the maximal expression rate of the bacteriocin (*KB*_*max*_) can be tuned through choice of promoter and ribosome binding-site [41]. The growth rates (*µ*_*max*_) can be tuned through choice of base strains or auxotrophic dependencies [42, 43]. The lower model posterior probability of *m*_66_ is reflected in the parameter density distributions, which are generally more constrained that those of *m*_62_. The distributions of *D* in both systems show a lower dilution rate is important for stable steady state. *KB*_*max*_ for all bacteriocins is tightly constrained to high maximal bacteriocin expression rates. For *m*_66_, the correlation coefficients for the two growth rates (*µ*_*max*1_, *µ*_*max*2_) show the parameters are loosely correlated. Additionally, we see that *N*_1_ has a higher maximal growth rate (*µ*_*max*1_) than that of *N*_2_ (*µ*_*max*2_). The faster maximal growth rate of *N*_1_ is necessary to counteract self-limitation that is negatively regulated by the population of *N*_2_. Conversely, *m*_62_ shows a wider distribution of strain growth rates at stable steady states and a low correlation coefficient, indicating these parameters are not as critical to produce the objective behaviour.

### Self-limiting motifs stabilise two strain systems

The dendrogram of Figure 2A highlights a cluster of high performing models that are closely related. This suggests underlying interactions of the model space exist that are important for producing communities with stable steady state.

Non negative matrix factorization (NMF) is an unsupervised machine learning method we can use to reduce the dimensionality of the interaction space [44]. We can use NMF to help us understand the underlying motifs and how they affect community stability. We represent each model by the interactions present in the system (Figure 2A). NMF takes these interactions and learns a number of clusters (*K*), models can be rebuilt by a weighted sum of these clusters. In our case, these clusters can be represented as interaction motifs. We set *K* = 4, in order to give us a digestible summary of the model space. Figure 3A shows the learned motifs that can be used to represent the entire model space. Figure 3B shows the component weights for each model, defining the membership each model has for each motif. The models are shown in descending order of posterior probability, we can see that *K* 2 is heavily weighted in the top performing models. The motif *K* 2 refers to self-limiting (*SL*) only interactions where the strain is sensitive to the bacteriocin it produces (Figure 3A, B). The top models are consistently assigned low weights for *K* 4 (Figure 3A, B), a motif which refers to other-limiting (*OL*) only interactions, where the strain is sensitive to a bacteriocin produced by the other strain (Figure 3A).

**Figure 3:**
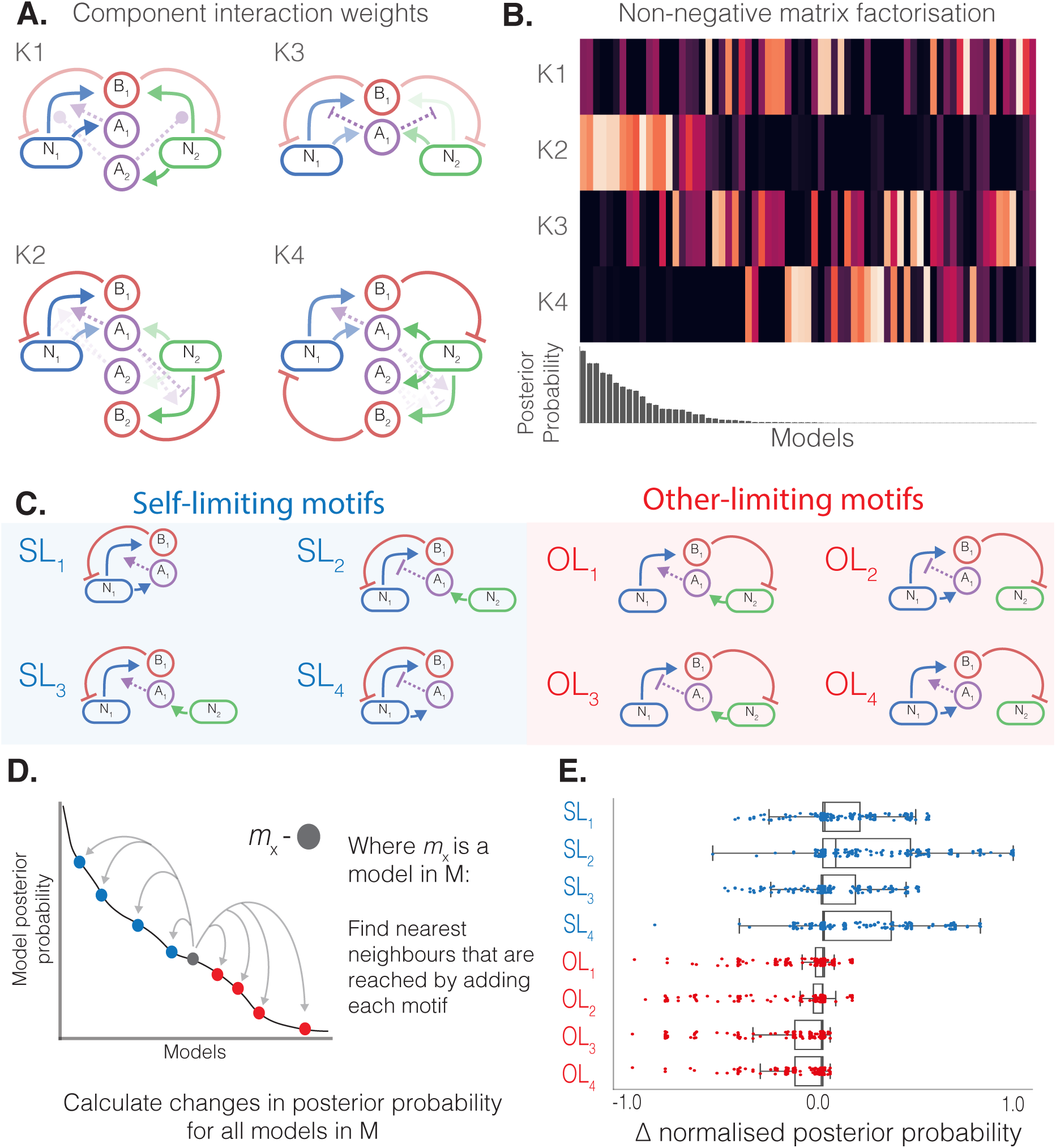
Contribution of network motifs to stability. A, B) Non negative matrix factorisation (NMF) analysis to learn motifs in model space using four components (K=4). A) Four components learned by NMF, the line opacity indicates the coefficient of the interaction. B) Visualisation of the component weights for each model. Each column is a model, with light colours corresponding to high weight and dark colours low weight. C, D, E) Manually curated minimal motifs capture interaction importance. C) Minimal motifs split into self-limiting (SL) and other-limiting (OL) by the direction of bacteriocin killing. D) Illustration of the algorithm used to generate each datapoint in E). Moving from a model to the nearest neighbours that can be built by adding a motif will produce a change in model posterior probability. E) Boxplots and scatter plot showing the change in posterior probability when adding each motif to a model. The boxplots show the median, first and third quartile. The lower and upper whiskers mark the 5th and 95th percentiles respectively.

The continuous learned motifs from NMF can be hard to interpret. We use the indications produced by NMF to curate our own discrete motifs. *K* 2 and *K* 4 show us the direction of bacteriocin sensitivity is an important feature and we proceed to investigate this further. All models can be built by combining eight fundamental motifs which can be categorised as either *SL* or *OL*, based on the direction of bacteriocin sensitivity (Figure 3C). Within each category, motifs are differentiated by the way bacteriocin expression is regulated (Figure 3C). For example, *m*_66_ = *SL*_2_, *m*_48_ = *SL*_4_ + *SL*_2_ and *m*_62_ = *SL*_2_ + *SL*_2_. In order to assess the importance of each motif for producing stable communities we perform a motif impact analysis (Figure 3D). For each model we identify the nearest neighbours in the model space that can be built by adding each motif. By calculating the change in posterior probability for each neighbour, we can summarise the contribution each motif brings to the model posterior probability (Figure 3D). By repeating this across the entire model space, we are able to quantify whether a motif is stabilising or destabilising (Figure 3E). The lower quartiles of *SL* motifs all show lower negative change magnitudes compared with the lower quartiles of *OL* motifs. The upper quartiles of *SL* motifs show a higher positive change magnitude than that of *OL* motifs. Together these show the addition of *SL* motifs more often result in an improved posterior probability, whereas addition of *OL* motifs are more often resulting in decreased posterior probability. The upper quartile of *SL*_2_ shows the motif has the most stabilising effect, closely followed by *SL*_4_. We see these findings are reflected by top models identified in Figure 2B, where all models are constructed with *SL*_2_ and *SL*_4_ motifs.

The total output of bacteriocin by a population will be a function of the population’s density. All *SL* motifs therefore possess a fundamental negative feedback relationship between growth rate and density, augmented by the mode of QS regulation. Conversely, the population density and growth rate of a strain in *OL* motifs are decoupled. This lack of feedback is a clear explanation as to why we see *SL* motifs as positive contributors to stability while *OL* motifs have a destabilising effect.

### Designing three strain communities that achieve steady state

While several studies have demonstrated the ability to establish synthetic two strain systems [45, 46, 23, 47, 48, 49, 20, 50, 51, 52, 53, 54], efforts with three strains are sparser [55, 24, 56]. Having demonstrated the automated design of two strain systems, we next tackle the far larger challenge of designing stable three strain communities. The addition of a single strain significantly increases the parameter space, engineering options and possible interactions. We define our available parts consisting of three strains (*N*_1_, *N*_2_, *N*_3_), three bacteriocins (*B*_1_, *B*_2_, *B*_3_) and two orthogonal QS systems (*A*_1_, *A*_2_). We maintain the same strain engineering restrictions, allowing up to one QS expression and up to one bacteriocin expression per strain. Each strain can be sensitive to up to one bacteriocin. Given the available parts and engineering limits, the Model Space Generator yields 4,182 unique models. Due to the much greater number of models, we group models based upon the interactions in each model by hierarchical clustering for up to five levels. The average posterior probabilities of each cluster are shown (Figure 4A). 3,289 models have a posterior probability of zero, highlighting how much more difficult this design scenario is. ABC SMC identifies *m*_4119_ as the system with the highest posterior probability for producing stable steady state. *m*_4119_ consists of a two QS molecules; *A*_1_ is produced by *N*_2_, *A*_2_ is produced by *N*_3_ (Figure 4B). The QS molecules repress the expression of self-limiting bacteriocins produced by each population. Using the minimal motifs defined in Figure 3C, *m*_4119_ can be summarised as *m*_4119_ = 3 × *SL*_2_. We subset the model space on the counts of heterologous expression in the system, yielding subsets containing candidate models with two, three and four expressed parts (low complexity to high complexity) (Figure 4B). Models with two heterologously expressed parts all had a posterior probability of 0.0 and are not shown. Again, we see a diminishing increase in posterior probability that comes with increasing complexity. *m*_3938_ is the more complicated neighbour of *m*_4119_, the difference being *N*_1_ is also contributing with production of *A*_1_, resulting in a fall in the posterior probability. The increase in posterior probability that occurs when moving from *m*_4125_ to *m*_4119_ has *BF* < 3.0, indicating the difference between the posterior probability of the two models is not substantial. These system comparisons highlight the trade-off between increasing complexity and improving system performance. In a similar fashion to the two strain model space, the top performing models are dominated by self-limiting only interactions (*Supplementary* Figure 1).

**Figure 4:**
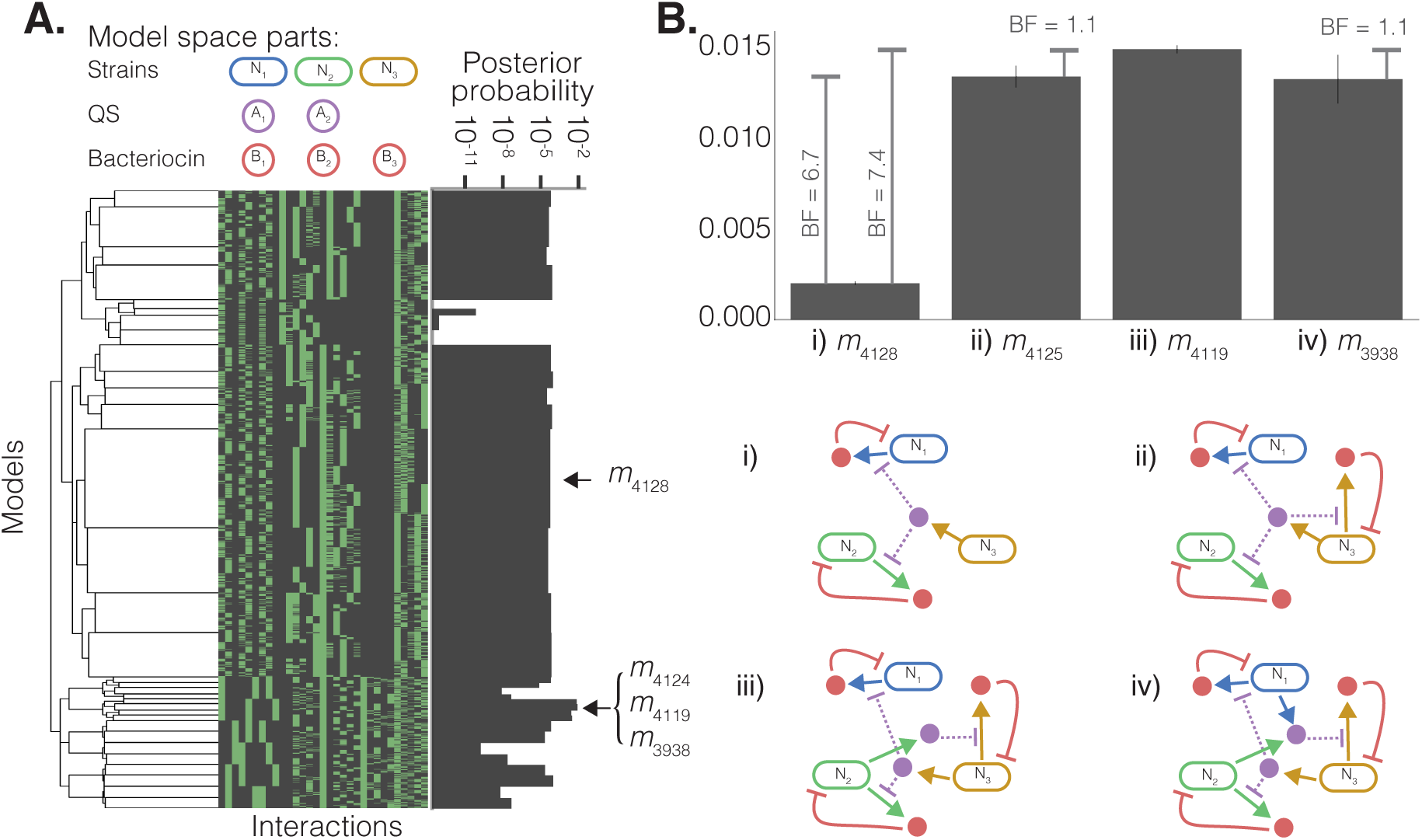
Output of AutoCD for three strain stable steady state objective A.) Dendrogram is generated by hierarchical clustering of the adjacency matrices for each model in the three strain model space. We set the limit of number of levels to 5, in order to show high level groups. Each column of the heatmap represents a possible interaction between species, where green indicates the interaction exists for the model and black indicates absence of the interaction. The posterior probability plot shows the average posterior probability within each group of models. B) Shows the models with highest posterior probability when subsetted for number of parts expressed, in order of increasing complexity (3, 4, 5 and 6 expressed parts). Bayes factors (BF) are shown for pairwise comparison of the three models and error bars show the standard deviation between three repeat experiments. Model schematics show the interactions between strains (blue, green and red), bacteriocin (red) and QS molecules (purple). Models with two parts showed posterior probability 0.0 and are not shown.

### Multiple engineered bacteriocins are more important than multiple orthogonal QS systems

Our results have identified top performing models in the two and three strain model spaces. We have also highlighted the diminishing returns that occur with increasing model complexity in top performing models. Next we aim to summarise the importance of different parts and their contribution to the stable steady state objective behaviour, further enabling us to triage genetic parts for construction in the lab.

Figure 5 shows a summary of the parts used to construct three strain systems and the average posterior probabilities they yield. This gives us important information to form heuristic rules in the design of three strain systems. Figure 5A shows the very similar posterior probability when comparing two QS systems rather than one. Figure 5B demonstrates the substantial advantage of repressive QS regulation of bacteriocin production over inducible systems. Figure 5C shows very strong evidence in favour of using three bacteriocins to produce stable steady state in three strain systems. These three statistics suggest that on average there is little advantage to be gained in the use of two QS systems, and priority should be given to the use of a single repressive QS to regulate three bacteriocin systems, such as we see in *m*_4125_.

**Figure 5:**
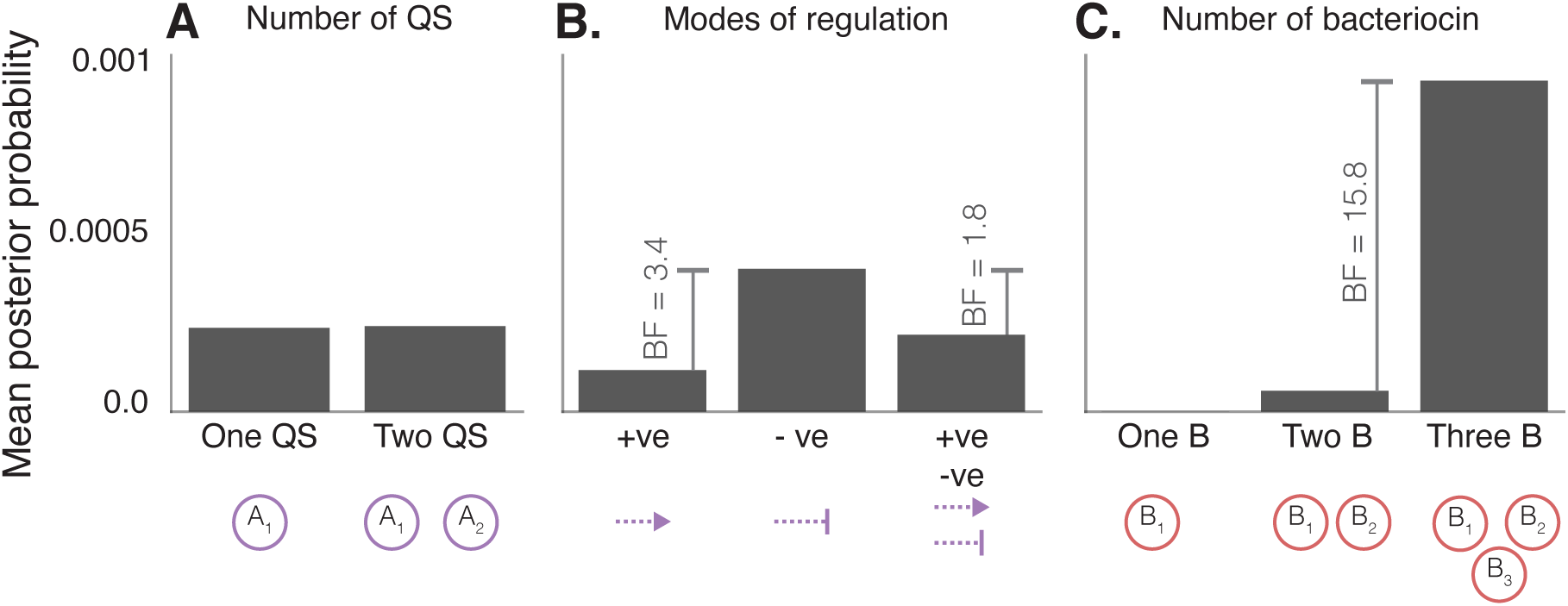
Summary of parts and by the mean posterior probabilities of subsets from three strain model space. A) Comparing systems with one and two orthogonal QS parts. B) Comparing modes of bacteriocin regulation in the system by subsetting for systems with induction (+ve), repression (-ve) or both (+ve, -ve). C) Comparing systems with one, two and three bacteriocin.

### Defining stable steady state population ratios in three strain systems

Synthetic communities can be applied to improving yields and efficiency via the distribution of bioproduction pathways between subpopulations [46, 53]. The density of a subpopulation will determine its productive output capacity for the function it conducts. Being able to define the steady state composition of a synthetic community is therefore an important feature. Here we demonstrate that secondary objectives can be applied to the output of ABC SMC without further simulation time, identifying key parameters that enable fine tuning of stable steady state population densities.

The 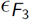 threshold value ensures all simulations in the final population have an *OD* > 0.001. Figure 6A shows the community composition distribution of m_4119_. The majority of accepted particles show a final community composition that is dominated by a single strain. By using the final population distances from ABC SMC we can subset for a secondary objective and identify how the system can be tuned to produce a more evenly distributed community composition. We set a secondary objective, stipulating that all strains must be of *OD* > 0.1 (pink) (Figure 6B). Therefore strains that do not meet the secondary objective have 0.001 < *OD* < 0.1 (blue) (Figure 6B). From these two subsets we generate separate parameter distributions and calculate the divergence using Kolmogorov-Smirnov (KS). Parameter distributions that show the greatest divergence are important for changing the system behaviour from one that is dominated by a single strain, to one that has a more even distribution of strain densities. The distributions of four parameters that exhibit greatest divergence are shown in Figure 6C. A higher dilution rate (*D*) and lower maximal bacteriocin expression rates 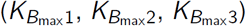 are associated with producing an more evenly distributed community composition. Importantly, all three parameters are realistically tunable. The dilution rate can be controlled directly through the chemostat device, while bacteriocin expression rates can be changed through the choice of promoters and ribosome binding sites.

**Figure 6:**
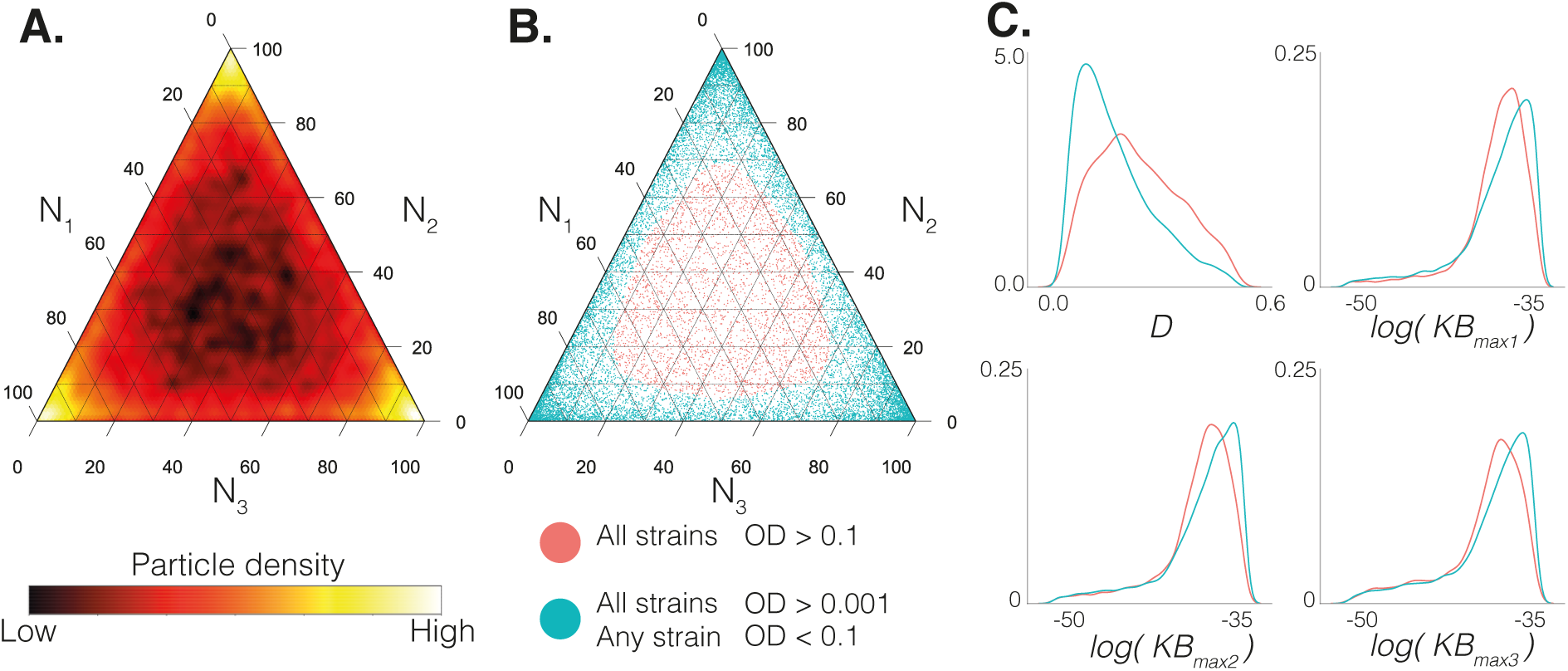
Distribution of population densities in Model 4119. Axes of ternary diagrams (A and B) show percentage composition of the community. A) Heat map showing the community composition at stable steady state in Model 4119. B) Scatter plot of the stable steady state systems, highlighting the secondary objective where all strains have *OD* > 0.1 (red), and the primary objective only where any strain has 0.001 < *OD* < 0.1. C) Density plots comparing the parameter distributions of four parameters that show the greatest divergence to produce the secondary objective: Dilution rate (*D*) and maximal bacteriocin expression rates (*KB*_max1_, *KB*_max2_, *KB*_max3_). The Kolmogorov-Smirnov values for the two objectives are 0.25, 0.12, 0.13 and 0.12 respectively.

## Discussion

Synthetic communities built to date have employed the use of QS, metabolic dependencies, intracellular lysis proteins, toxins and extracellular AMPs to engineer interactions that enable community formation [55, 54, 24]. Studies often incorporate mathematical modelling to inform experimental design and identify the conditions that will produce the expected population dynamic. The design of the fundamental interactions of the system itself is often directed by mimicking ecological interactions found in nature, or by rational judgement. As the possible types of engineered interaction increases, so does the need for comprehensive assessment of the vast model spaces. The modelling and statistical framework demonstrated here addresses this design problem. With our examples we have highlighted important design features and heuristic rules for building synthetic steady state communities. Comparative coculture studies have been used to identify stable partnerships in microbial ecosystems [57, 58]. These methods could be used to validate our findings in the wet lab by constructing a subset of strains from the model space and comparing the ability of pairs to produce stable steady state.

We have identified optimal system designs using bacteriocins and QS for stable steady state in two and three strain communities. *m*_62_, the top model of the two strain model space uses a cross-protection mutualism, whereby the density of each subpopulation inhibits the self-limitation of the other. Cross-protection mutualism has previously been incorporated in synthetic microbial communities via the mutual degradation of externally supplied antibiotics [49]. Metabolic interdependencies can also be employed to engineer mutualism [50, 51]. Similarly, in *m*_4119_ of the three strain model space (Figure 4B) we see pair-wise cross-protection mutualism between two subpopulations and a dependent subpopulation. All top performing models used SL interactions to produce stable steady state dynamics. Self-limitation is observed in many natural biological communities, normally in a response to stress [59, 60]. These processes, while detrimental to the individual, provide a net benefit to the community through release of a public good - they are altruistic processes [61]. Altruistic cell death is conserved throughout different species implying a competitive advantages in natural environments [62]. SL interactions have previously been employed for building synthetic co-cultures. Scott *et al.* demonstrated the use of QS regulated lysis protein expression to implement a self-limiting interactions in two strain systems, overcoming competitive exclusion in a batch culture.

The robustness of SL interactions can be explained by the feedback loops involved. Total bacteriocin output by a subpopulation is heavily dependent upon its population density; low population density will naturally have a low output of bacteriocin [63], making QS a secondary level of regulation. This is supported by both two and three strain scenarios where we observe the diminishing returns that come with increasing complexity. Figure 5 shows that increasing the number of bacteriocin in a system yields greater increases in stability than increasing the number of QS systems. A closed feedback loop exists between the bacteriocin expression rate and the population density. This is an important reason why we see all *SL* motifs generally show positive contribution to stability. Conversely, in *OL* motifs the population expressing the bacteriocin will not be negatively affected and therefore a closed feedback-loop does not exist. It should be emphasised that while the design rules we have identified hold true for a stable steady state objective, it may not be the case for other objective population dynamics, such as oscillations. However, new objectives can be investigated by changing the distance functions which describe the population dynamic.

Finally, we showed that the posterior parameter distribution from ABC SMC can be used to make decisions on part characteristics and experimental conditions (Figure 2C and Figure 6C). Our results show the dilution rate (*D*) is an important experimental parameter for producing stable steady state, and tuning the community composition. The rate of removal of molecules from the environment can produce very different population dynamics. This result is supported by previously conducted work where the dilution rate has regularly been demonstrated as important for determining the population dynamic [23, 49, 6]. We also show our methodology can identify systems that are robust to differences in growth rate, highlighted by the comparison of *m*_66_ and *m*_62_ in Figure 2C. Together these draw attention to important part characteristics that should be considered when constructing a stable community. Furthermore, it should be emphasised that if the parts are already selected and characterised, their parameters can be fixed in the prior distribution reflecting the known behaviour. The framework we have developed offers a natural entry point to the design-build-test cycle, providing a data informed roadmap for building a robust synthetic community with a desired behaviour. We have revealed stable steady state systems in a two and three strain model space, and generated impactful rules and heuristics for their construction. The flexibility of this framework enables us to quickly redefine population level behaviours depending on the required application.

## Methods

### Model space generator

Models are generated from a set of parts which are expressed by different strains in the system. We represent an expression configuration through a set of options. We define the options for expression of *A* in each strain, where the options are not expressed, expression of *A*_1_, and expression of *A*_2_ (0, 1 and 2). We define the options for expression of bacteriocin, which for the two strain model space includes no expression, expression of *B*_1_ or expression of *B*_2_ (0, 1, and 2). For the three strain model space, this includes includes no expression, expression of *B*_1_, expression of *B*_2_ or expression of *B*_3_ (0, 1, 2 and 3 respectively). Lastly we define the mode of regulation for the bacteriocin, which can be either induced or repressed (0 and 1). This is redundant if a bacteriocin is not expressed.

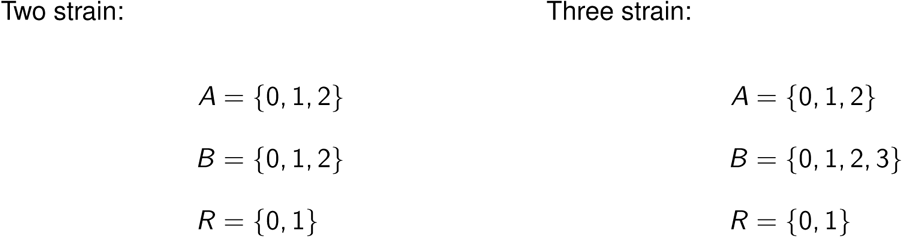

This enables us to build possible part combinations that can be expressed by a population. Let *P*_*c*_ be a family of sets, where each set is a unique combination of parts.

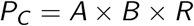

Each strain in a system can be sensitive to up to one bacteriocin. Let *I* represent the options for strain sensitivity. In the two strain model space, the options are insensitive, sensitive to *B*_1_ or sensitive to *B*_2_ (0, 1, and2 respectively). In the three strain model space, where the options are insensitive, sensitive to *B*_1_, sensitive to *B*_2_ or sensitive to *B*_3_ (0, 1, 2 and 3 respectively).

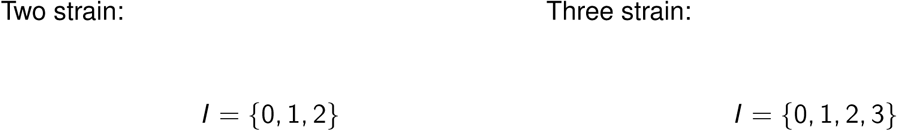

Each strain is defined by it’s sensitivities, and expression of parts. Let *P*_*E*_ be all unique engineered strains:

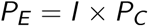

Which can be combined to form a model, yielding unique combinations in two strains and three strains:

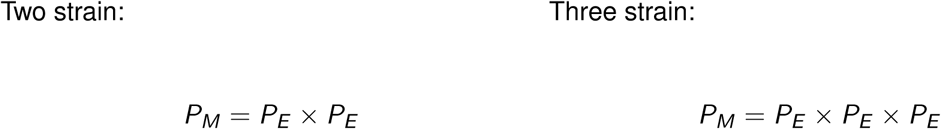

Finally, we use a series of rules to remove redundant models. A system is removed if:

1. Two or more strains are identical, concerning bacteriocin sensitivity and combination of expressed parts.
2. The QS regulating a bacteriocin is not present in the system.
3. A strain is sensitive to a bacteriocin that does not exist in the system.
4. A bacteriocin exists that no strain is sensitive to.

This cleanup yields the options which are used to generate ODE equations for system.

### System equations

s in each system are rescaled to improve speed of obtaining numerical approximations.

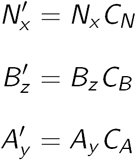

Each model is represented as sets of species

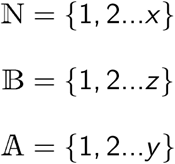

Species are represented as differential equations

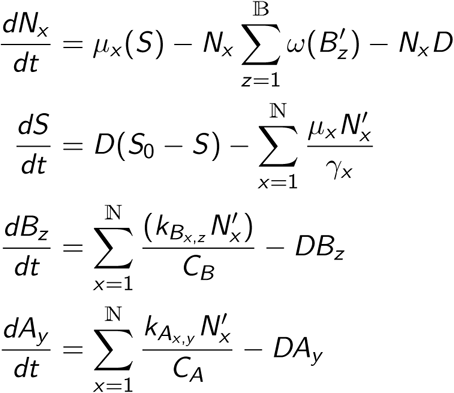

Growth modelled by Monod’s equation for growth limiting nutrient (*S*)

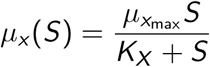

Killing by bacteriocin, where *ω*_*max*_ = 0 if strain is insensitive

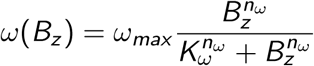

Induction or repression of bacteriocin expression by QS species, *A*_*y*_

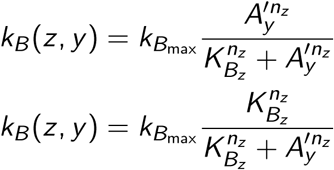

Simulations were conducted for 1000hrs, the final 100hrs were to calculate the summary statistics and were stopped early if the population of any strain fell below 1*e*−10 (extinction event). Simulations with an extinction event have distances set to maximum. This was done to prevent excessive time spent solving ODEs of collapsed populations.

### Bayesian inference

Let *θ* ∈ Θ be a sampled parameter vector with a prior *π*(*θ*). Given an objective of *x*_0_, where *x*_0_ exists in the solution space, *x*_0_ ∈ 𝒟. We define the likelihood function for the objective behaviour as *f* (*x*_0_|*θ*). Bayes’ theorem gives us the posterior distribution of *θ* that exists for the objective *x*_0_.

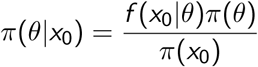

We can rewrite *π*(*x*_0_) where *a* and *b* represent the lower and upper bounds of the parameter value:

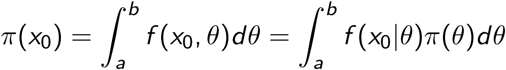

The posterior distribution informs us of the parameter distribution that gives rise to the objective.

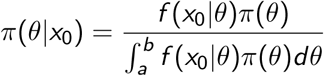

Let *m* be a model from a vector of competing models, *M*, such that *m* ∈ *M* = {*m*_0_, *m*_2_…*m*_*n*_}. Each model has its own parameter space, allowing us to define a joint space, (*m, θ*) ∈ *M ×* Θ_*M*_.

We can write Bayes’ theorem in the context of a model space.

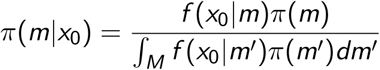

Since the *M* is discrete, we can rewrite this as

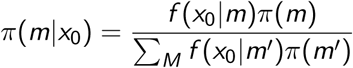

The marginal likelihood of the model, *f* (*x*_0_|*m*), considers the joint parameter and model space

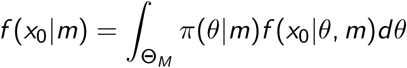

### Approximate Bayesian computation

Writing the likelihood function, *f* (*x*_0_|*θ*), in terms of summary statistics can be difficult. We bypass this and approximate the posterior by generating data from a model. We can sample a parameter vector from the prior, *θ*^*^ ∼ *π*(*θ*), which is simulated to yield a data vector, *x*^*^. This can be written as a conditional, *x*^*^ ∼ *f* (*x*|*θ*^*^), which also gives the joint density, *π*(*θ, x*).

In order to obtain the posterior distribution that satisfies our objective behaviour, *x*_0_, we apply a conditional to define whether a generated data vector, *x*^*^ belongs to the objective *x*_0_.

If *x* = *x*_0_

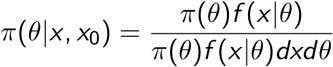

Else

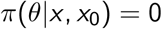

Let *ρ*(*x, x*_0_) be a distance function that compares a simulation to the objective. Using distance thresholds (*ϵ*), we can define values below which the distance is acceptably small. We can redefine *π*(*θ*|*x, x*_0_) in the context of thresholds to obtain an approximation of the posterior.

If *ρ*(*x, x*_0_) < *ϵ*

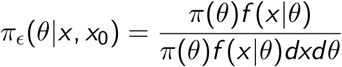

Else

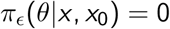

The smaller *ϵ* is and the larger the number of simulations conducted, the more accurate the representation of the true posterior will be. We can write this marginal posterior distribution as

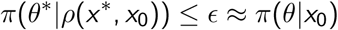

### Model selection with ABC SMC

The most basic ABC algorithm is the ABC rejection algorithm. Let *ϵ* be the distance threshold the defining the necessary level of agreement between the objective, *x*_0_, and a given simulation, *x*^*^. In this paper we use a variant of ABC, ABC Sequential Monte Carlo (ABC SMC) [39]. Particles are sampled from the prior distributions, each particle represents a sampled model and sampled parameters for that model. ABC SMC evolves particles sampled from the prior distribution through a series of intermediate distributions. Particles sampled from an intermediate distribution are perturbed and and given an importance weighting to define their sample probability for the next distribution. The distance threshold (*ϵ*) is decreased between distributions, moving the acceptance criteria closer to the objective. These features aim to improve the acceptance rate of particles while maintaining a good approximation of the posterior distribution.

#### Algorithm 1: Model selection with ABC SMC

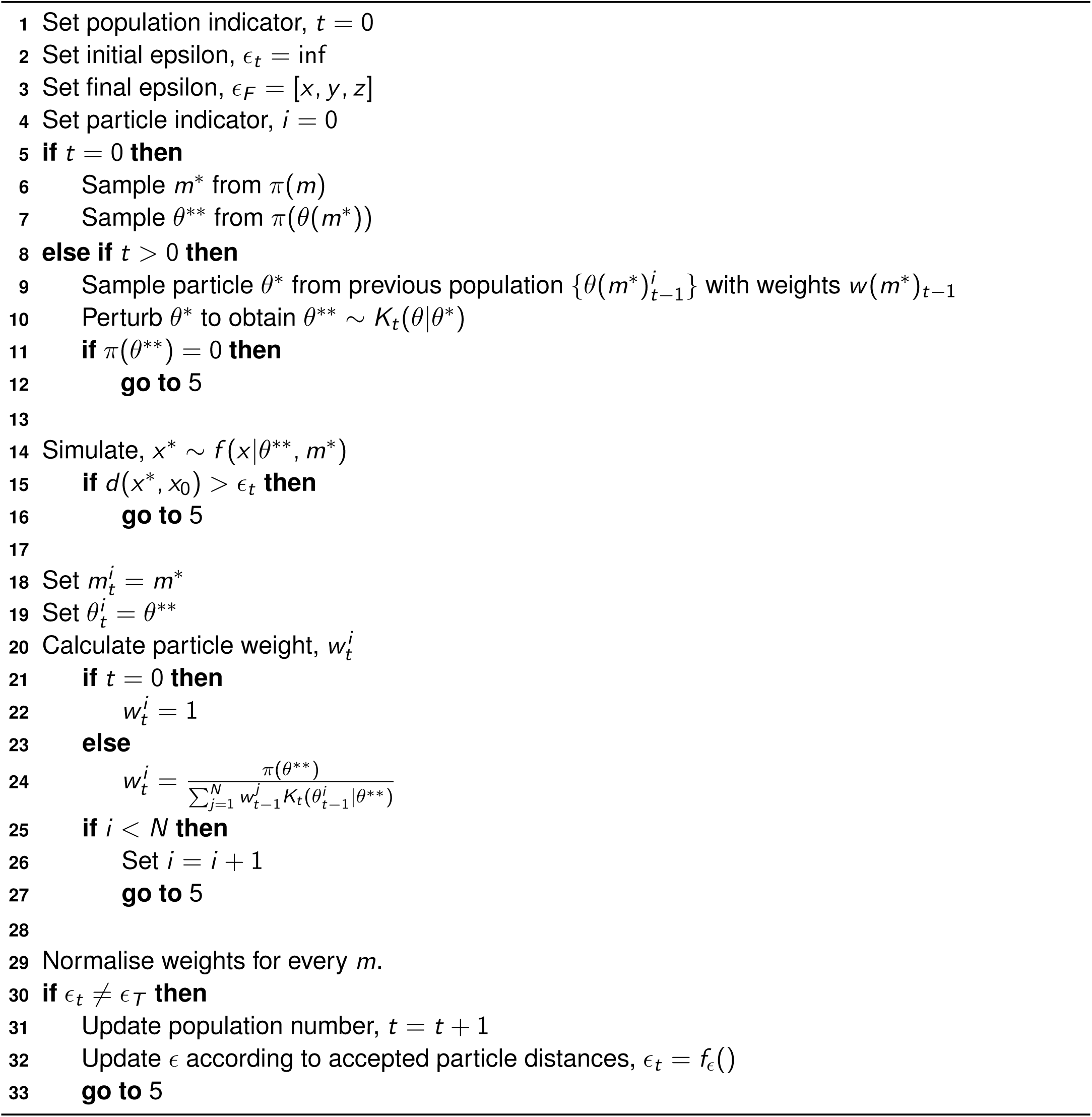

The Bayes factor can be used to help us interpret how much better (or worse) one model is than the other. Given two models, *m*_1_ and *m*_2_, the Bayes factor is calculated as:

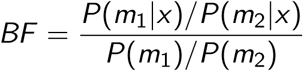

*P*(*m*_*i*_) is the prior, and *P*(*m*_*i*_ |*x*) is the posterior probability. Given uniform priors, *P*(*m*_*i*_) = 1*/M*, where M is the number of models. Therefore we can simplify to:

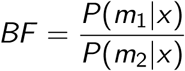

The Bayes factor is a measure the support for *m*_2_ relative to *m*_1_. It accounts for the number of parameters, or complexity of the two models. The Bayes factor allows us to directly compare the weight of evidence for and against the two models and has the advantage that it can be used to compare non nested models. Two Bayes factors can be compared directly, since they both represent evidence in favour of the hypothesis [39, 40]. We therefore use Bayes factors to directly compare the ability of two models to represent the objective population behaviour. The following table allows us to interpret *BF*.

### Software packages and simulation settings

Repository of AutoCD can be found at https://github.com/ucl-cssb/AutoCD/. The repository includes configuration files for the two- and three- strain experiments. ABC SMC model selection algorithm was written in python using Numpy, Pandas and Scipy. ODE simulations were conducted in C++ with a Rosenbrock 4 stepper from the Boost library. All simulations use an absolute error tolerance of 1*e*−9, and relative error tolerance of 1*e* − 4. Non negative matrix factorisation was conducted using Scikit Learn. Dendrograms were made from Scipy, using the unweighted pair group method with arithmetic mean (UPGMA) clustering algorithm. Ternary diagrams were made using python package python-ternary [64]. Parameter distribution plots were made in R using ggplot2.

### Prior distributions

Prior distributions for both two and three strain systems are sampled from uniformly between the min and max values listed below. Constant parameters have the same min and max value.

