## Supplementary Figures for "Automated design of synthetic microbial communities"

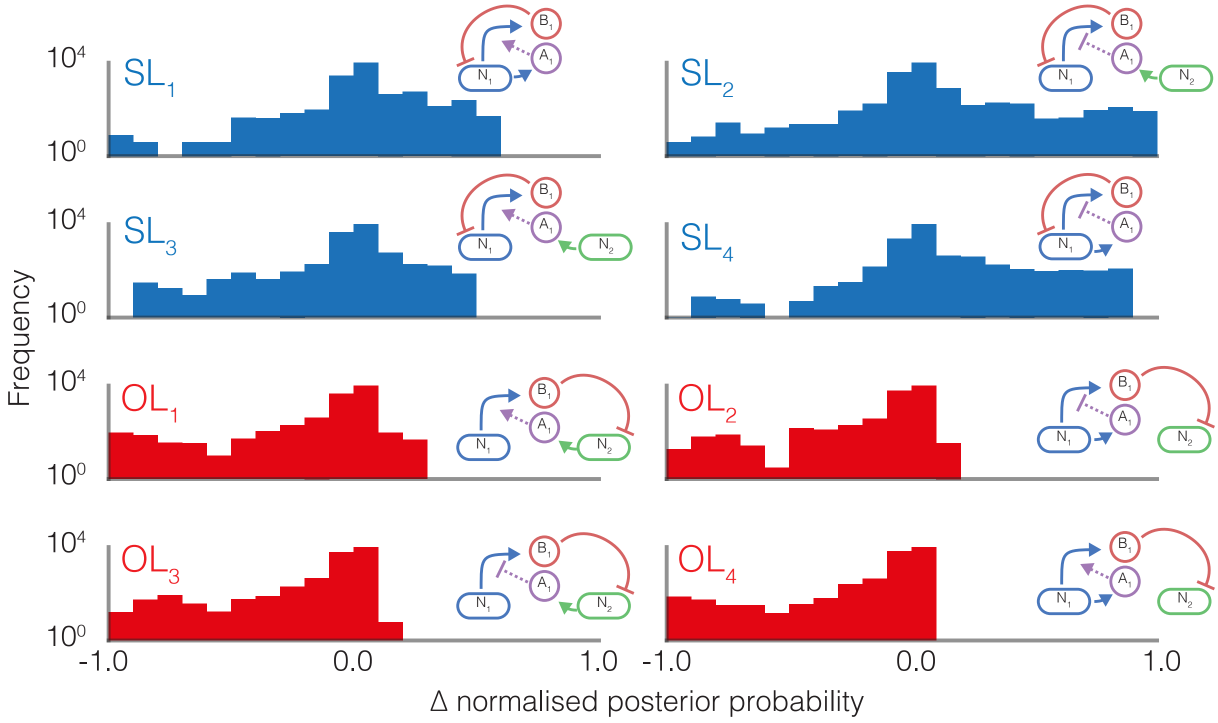


Figure 1: Histograms showing the change in posterior probability when adding each motif to a model in the three strain model space.
